# WaveletSEG: Automatic wavelet-based 3D nuclei segmentation and analysis for multicellular embryo quantification

**DOI:** 10.1101/2020.07.24.220285

**Authors:** Tzu-Ching Wu, Xu Wang, Linlin Li, Ye Bu, David M. Umulis

**Affiliations:** Department of Agriculture and Biological Engineering, Purdue University, West Lafayette, IN 47907, USA; Weldon School of Biomedical Engineering, Purdue University, West Lafayette, IN 47907, USA

## Abstract

Identification of individual cells in tissues, organs, and in various developing systems is a well-studied problem because it is an essential part of objectively analyzing quantitative images in numerous biological contexts. We developed a size-dependent wavelet-based segmentation method that provides robust segmentation without any preprocessing, filtering or fine-tuning steps, and is robust to the signal-to-noise ratio (SNR). The wavelet-based method achieves robust segmentation results with respect to True Positive rate, Precision, and segmentation accuracy compared with other commonly used methods. We applied the segmentation program to zebrafish embryonic development IN TOTO for nuclei segmentation, image registration, and nuclei shape analysis. These new approaches to segmentation provide a means to carry out quantitative patterning analysis with single-cell precision throughout three dimensional tissues and embryos and they have a high tolerance for non-uniform and noisy image data sets.

## Introduction

In developmental biology, quantification of cell morphology, tissue patterning, and gene expression within a developing embryo at different stages provides detailed information that drives our understanding of differentiation and the function of signaling networks driving cell fate specification. Several nuclei segmentation methods have been developed to track nuclei position in fluorescent images and extract the spatiotemporal expression of each nucleus(McMahon *et al*., 2008)(Keller *et al*., 2008)(Stegmaier *et al*., 2014)(Meilhac *et al*., 2009). Most of the nuclei segmentation algorithms need image pre-processing steps before segmentation, and the settings in pre-processing are highly dependent on the microscopy imaging environment(Win *et al*., 2018). Global and adaptive thresholding segmentation methods are the simplest and the most common nuclei segmentation methods that use single or multiple threshold values on an image histogram to distinguish between an object and background(Otsu, 1979)(Cai *et al*., 2014).

However, most real images lack significant thresholding points or low contrast foreground and background(Vala and Baxi, 2013). The Watershed method is another common segmentation method also capable of dividing overlapping nuclei(Vincent, Vincent and Soille, 1991)(Beucher, 1992). Nearby pixels with similar features are classified topologically and defined as a catchment basin that are separated by a watershed ridge line from the adjacent catchment basin. Using the watershed method can divide overlapping objects, however, may cause an over-cutting problem. Other methods such as active contour models (ACMs)(Dufour *et al*., 2005)(Zhang, Zimmer and Olivo-Marin, 2004), level-set(Dzyubachyk *et al*., 2010) and graph cuts(Al-Kofahi *et al*., 2010) are also applied in the field of nuclei segmentation. Various level-set and model-based methods have been developed for accurate embryo nuclei segmentation(Santella *et al*., 2010)(Gudla *et al*., 2008)(Ambühl *et al*., 2012), although they are computationally expensive or require pre-knowledge(Irshad *et al*., 2014). To segment and track chromosome dynamics in the early *Drosophila* embryo, a fast level-set method was proposed(Yau and Wakefield, 2007). A 3D level-set method was also applied to detect nuclei and mitotic chromosomes in *Drosophila* embryos(Chinta and Wasser, 2012). A region-based active contour method was developed in 3D cell counting and segmentation of vertebrae in early embryogenesis(Rizzi and Sarti, 2009). We applied a number of these methods to quantify nuclei number and shape in zebrafish, however we encountered a number of problems associated with imaging large amounts of nuclei in a relatively large sample. Specifically, intensity attenuation effects due to absorption and scattering of fluorescent light in deep layer specimens can lead to intensity decay and underestimates in the segmentation results(Xu *et al*., 2015). Moreover, densely packed nuclei are often present in an embryo which requires overlapping nuclei detection strategies(Toyoshima *et al*., 2016). To develop the new approaches, we leverage wavelet transforms that are common in signal processing and are applied in many fields where the method is used to isolate the object from a noisy signal. The 2D WTMM (2D wavelet transform modulus maxima) method was first introduced for multiscale edge detection and segmentation of regions of interest in cell nuclei(Arnéodo, Decoster and Roux, 2000)(Arnéodo *et al*., 2003) and CT detection(Branco and Pombo, 2006).

Several image analysis software packages have been established, such as ImageJ(Abràmoff, Magalhães and Ram, 2004), Fiji(Schindelin *et al*., 2012), and instrument-bundled packages: ZEN and cellSens. However, commercial bundled packages lack flexibility and may lack affordable options for on-the-fly segmentation, and open source tools are either limited in specific applications, not robust in segmentation results, or not convenient to use. For nuclei segmentation tools, widely used software CellProfiler(Carpenter *et al*., 2006)(McQuin *et al*., 2018), provides pre-processing, object identification, and cell counting modules, but it is limited to only 2D images. Additional related software includes Matlab toolbox OML that is capable of automatic cell segmentation. Spatzcells(Skinner *et al*., 2013), FISH-quant(Mueller *et al*., 2013) and MINS(Lou *et al*., 2014) are Matlab executable codes that can generate segmentation cell masks and spot recognition for fluorescent 2D or 3D images. LSDCAS (The Large-Scale Digital Cell Analysis System)(Ianzini *et al*., 2002)(Davis *et al*., 2007) provides the ability for live cell image segmentation and tracking of cell trajectories. RACE(Stegmaier *et al*., 2016) was designed for automated 3D cell segmentation and cell reconstruction for embryos, based on a watershed segmentation approach that, in our hands is more sensitive to non-uniform or variable-quality images (Table S1).

In this paper we present a wavelet-based automatic 3D nuclei segmentation method that is object size-dependent, robust to noise and intensity attenuation, and can divide overlapping nuclei (Fig. S1-3). It contains six steps, including a 2D continuous wavelet transform, multi-scale object identification, 3D alignment, first and second nuclei division steps and outlier removal.

Unlike many segmentation methods, the wavelet-based method has a strong ability to segment objects in noisy images without any preprocessing steps. The method herein combines a 2D continuous wavelet transform (Fig. 1), and the only parameter needed for object determination is the wavelet scale factor determined by target object size, and no other preprocessing steps are needed. Through comparison between several wavelet coefficient maps, wavelet-based segmentation can easily distinguish the object region from the background in images and it can do this without any preprocessing or thresholding steps. We tested the model against manually curated 3D embryonic data and found that it has robust and stable segmentation performance with respect to True Positive Rate (TP rate), precision, and segmentation accuracy when compared to other methods. We integrated the wavelet-based nuclei segmentation, image registration, topology and geometry packages into an automatic interactive analysis platform named WaveletSEG for embryological developmental studies. Additionally, we developed a 3D ground truth annotation tool, synthetic image generator, and a segmented training dataset export tool and data visualization interface for additional data analysis and data validation in WaveletSEG. We quantified 3D nuclei and intensity spatial distributions, topology features and shape classification results at 4.7, 5.7, and 6.5 hours post fertilization (hpf) zebrafish using WaveletSEG.

**Fig. 1.**
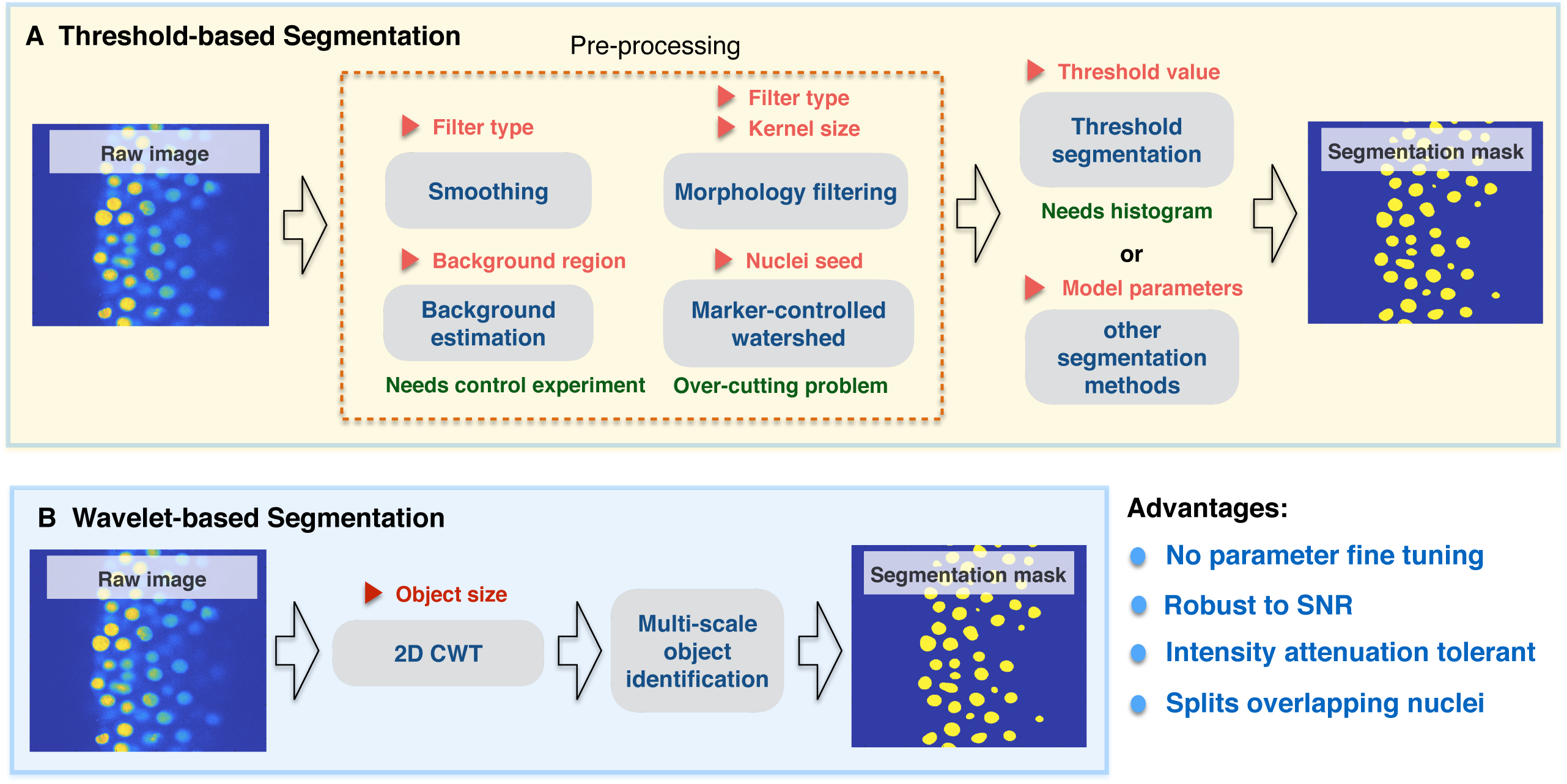
Comparison between threshold-based segmentation and wavelet-based segmentation methods. Preprocessing steps are crucial for most threshold-based segmentation methods and determine the quality of segmentation performance. Increased parameter numbers in preprocessing steps and segmentation method increases result complexity. Red triangle symbols indicate parameters required for each step and may need parameter fine tuning.

## Results

### Main steps of the wavelet-based segmentation method

The six main steps of the wavelet-based nuclei segmentation method (Fig. 2) are: (1) application of a 2D continuous wavelet transform (2D CWT), (2) multi-scale object identification, (3) 3D object alignment, (4) first round of nuclei division based on xy plane wavelet coefficients, (5) second division of nuclei based on z direction, and (6) deletion of outliers and small objects. In step one, a two-dimensional continuous wavelet transform is applied to each Z-stack slice of 3D sections common as output from confocal imaging (see Online Methods and Fig. 2B). While a number of wavelet functions were tested, we find that the 2D Mexican hat function as the mother wavelet function provides the ideal function to resolve point wise or nuclear-sized structures. The wavelet scale factor range applied after the transform is then based on the object size we wanted to identify. In line with this, we designed an efficient multi-scale search algorithm that compares different scales of the CWT coefficients and groups shape-similar regions such as the nuclei masks based on contour line similarities in the CWT coefficients diagram (see Online Methods, Fig. 2C and Fig. S4). Through the comparison between wavelet coefficient matrices over different scales we determined that a similarity index of 0.9 for between contour line similarity routinely identified nuclei without additional prepossessing or thresholding steps. Regions that meet the criteria are masked and labeled as nuclei in each stack for further processing to identify nuclei properties including progression through the cell cycle.

**Fig. 2.**
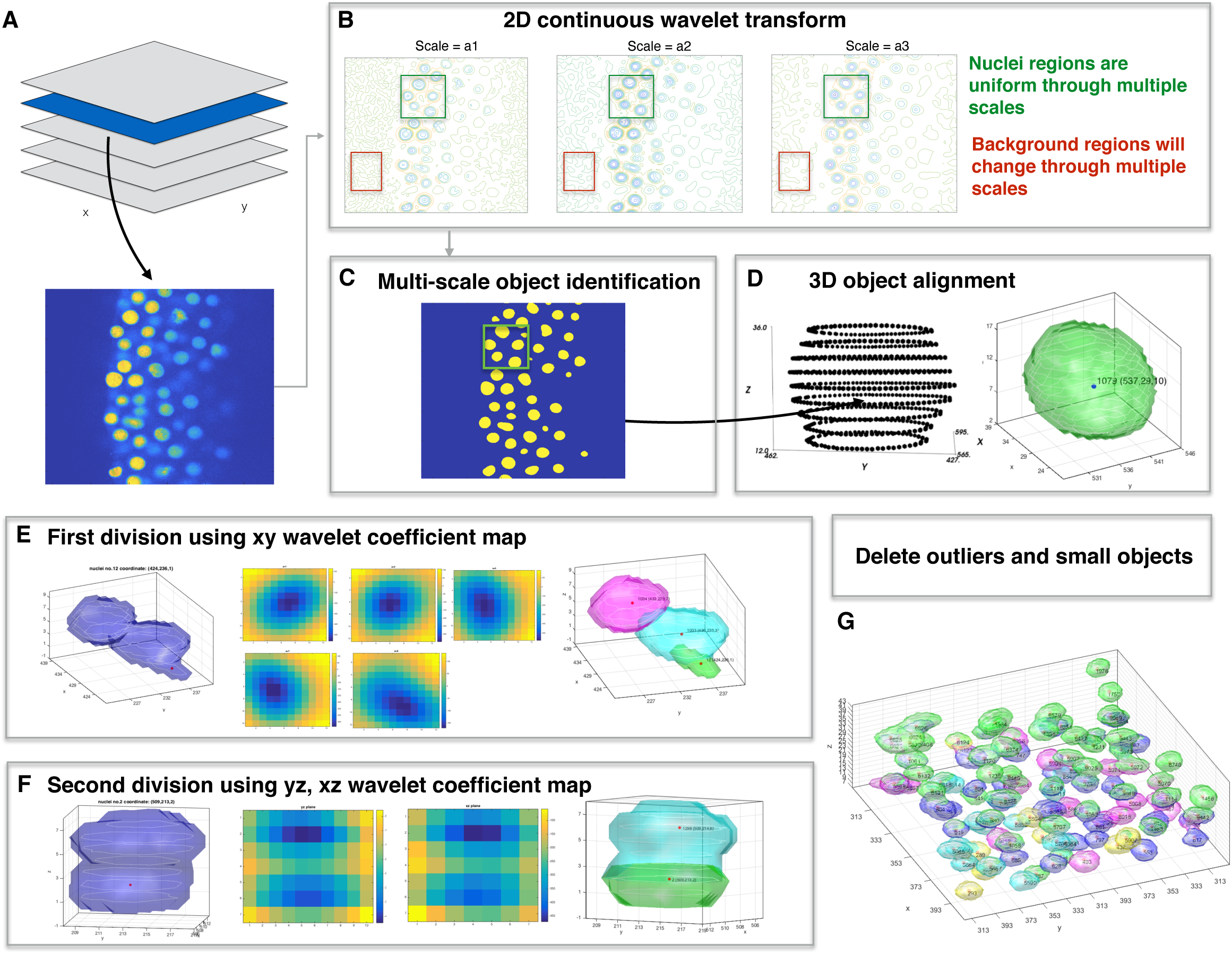
Overview of wavelet-based nuclei segmentation method. (A) Import 3D raw image files. (B) After importing 3D images, 2D continuous wavelet transform with appropriate wavelet scale factors are applying to each 2D z-stack. (C) 2D segmentation masks are obtained by applying multi-scale object identification to wavelet coefficient maps from previous step. (D) In 3D object alignment step, connected identified 2D masks from neighboring slices forms a 3D foreground object. (E) First division step if more than one center or center shift on neighboring z slice wavelet coefficient maps. (F) Second division step in z direction if multiple centers found on yz or xz wavelet coefficient maps. (G) Delete 3D nuclei voxels smaller then nuclei size × 0.2.

Following 2D identification, nuclei are reconstructed by 3D object alignment that aligns 2D nuclei cross-sections to recreate 3D nuclei shapes by comparing consecutive nuclei segmentation results from the preceding step (see Online Methods and Fig. 2D). The method computes the intersection of the mask coverage area of the neighboring z-plane. If the intersection area among adjacent z-planes is greater than 80% and if the max intensity is not lower than 20% nuclei center intensity, it is inferred that the adjacent mask coverage area should belong to the same 3D nuclei object.

In threshold and other segmentation approaches, a major challenge is errors in nuclei identification due to imaging conditions that lead to irregular or poor contrast boundaries or intensity attenuation that systematically distort measured fluorescence intensity. These factors compound in 3D and additional steps are needed for identification. First, the wavelet coefficient local minimum on each z-plane is determined because the spatial center of the object will produce a local minimum on the wavelet coefficient z-plane. If there is a shift in the wavelet coefficient local minimum on two consecutive z-planes and the shift distance is larger than 1/3 nuclei radius, this is indicative of two different nuclei objects, and it will be marked as likely belonging to different nuclei (see Online Methods, Fig. 2E, and Fig. S5). For identification of potential separate overlapping nuclei in the YZ-plane and XZ-plane a similar center point detection is used (Fig. 2F, and Fig. S6). If multiple wavelet coefficient local minimum points are found on both the YZ-plane and the XZ-plane it is likely they come from two nearby nuclei (see Online Methods). Lastly, noise or false positive nuclei after the additional segmentation are removed based on object size filtering (Fig. 2G).

### Evaluation of segmentation methods with synthetic and real images

To benchmark the wavelet and other methods that are coded into the WaveletSEG program, we developed tunable test data as well as a ground-truth data to manually segment 3D images for validation. In addition to the wavelet method, the following methods are included in the program: point wise method(Zinski *et al*., 2017), Otsu method(Otsu, 1979), and Derivatives Sum (DS) method(Rajasekaran *et al*., 2016). The validation datasets we used to evaluate segmentation performance included synthetic image datasets that were created by using a synthetic data generator (see Online Methods and Fig. S7), publicly open image datasets(Ulman *et al*., 2017), and our ground truth image dataset determined by multiple rounds of manual segmentation. We also generated synthetic nuclei overlapping datasets to test the ability to divide overlapping nuclei.

Three evaluation criteria were applied to compare the accuracy of four segmentation methods. If a representative point was nearest-neighbor of a point in ground truth dataset and visa-versa, the object was regarded as a true positive (TP). If only the former condition was met, the segmented result is considered a false negative (FN). If only the latter condition was met, the object was regarded as a false positive (FP). Then the true positive rate (TP rate) is equal to the true positive number divided by the ground truth number, and Precision is defined as the TP number divided by the sum of TP number with FP number. TP rate defined in equation (1) represents the ratio of successful nuclei segmentation within all ground truth nuclei data, and Precision is defined in equation (2). The false negative rate is the false negative number divided by the ground truth number. We also use segmentation accuracy to estimate pixel-wise segmentation performance, which is defined as the overlapping region between ground truth segmentation region and segmentation region in pixel level.

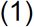

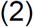

We use the Jaccard similarity index J (equation (3)) to define segmentation accuracy^39^ which means the amount of overlap between the segmentation results S and ground truth annotation results R.

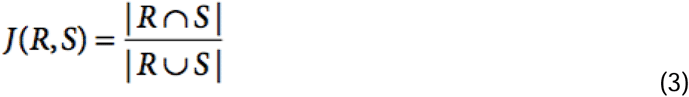

In test data we added multiple different types of image noise that would normally be encountered in an imaging environment. For Gaussian white noise (Fig. 3A), the wavelet-based segmentation method has the best TP rate and Precision followed by DS method. Otsu method obtained the lowest Precision and the point-wise method performed the poorest in TP rate. In salt and pepper noise cases (Fig. 3B), both TP rate and Precision were equal to one when segmented by the wavelet-based method, and all other method results are lower than 0.9 in both TP rate and Precision. In images with Intensity attenuation (Fig. 3C), the point-wise method has very high false positive rates and Otsu method performed the poorest in TP rate. In this test, the wavelet-based segmentation method had the highest TP rate and Precision. In the next step we examined the ability to divide nuclei in overlapping images for four segmentation methods (Fig. 3D). Compared to three other segmentation methods, the wavelet-based method has the best ability to divide overlapping nuclei in both TP rate and Precision.

**Fig. 3.**
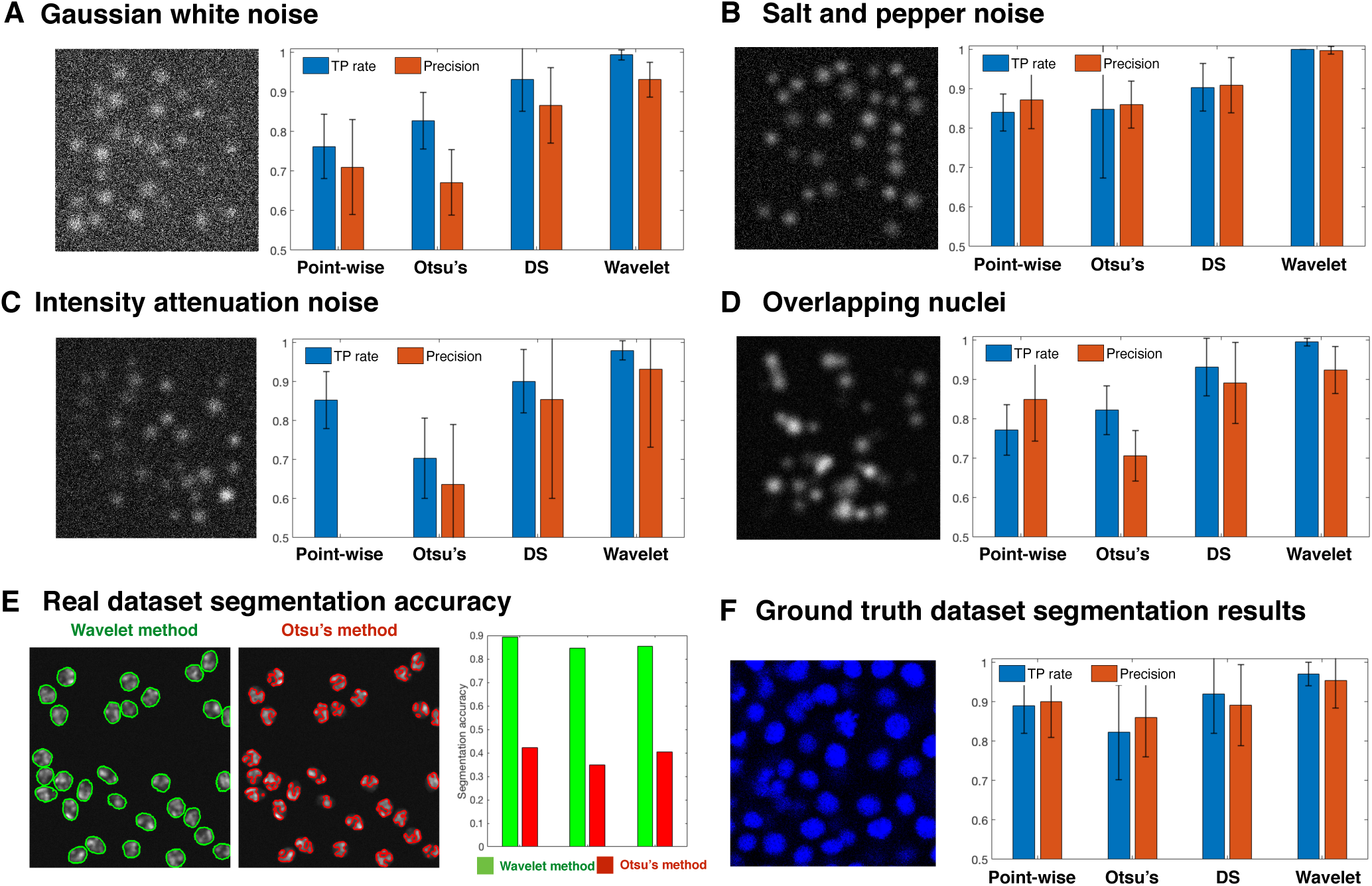
Performance criteria and segmentation accuracy of four segmentation methods on synthetic images and ground truth image datasets. (A-D) True positive (TP) rate and Precision on four kinds of noisy synthetic images using three segmentation methods and the wavelet method. Error bars indicate standard deviation. (A) Samples of synthetic images (N=20) Gaussian white noise and TP rate and Precision when applying point-wise method, Otsu method, DS method and wavelet-based segmentation method. (B) Samples of salt and pepper noise synthetic images (N=20) and TP rate and Precision for four segmentation methods. (C) Intensity attenuation noisy synthetic image samples (N=20) and TP rate and Precision for four segmentation methods. (D) Samples of overlapping synthetic image samples (N=20) and TP rate and Precision for four segmentation methods. (E) Segmentation accuracy when applying Otsu method and wavelet-based segmentation method on three Fluo-N2DH-GOWT1 sample images. (F) True positive rate and Precision on five ground truth images when using point-wise method, Otsu method, DS method and wavelet-based segmentation method.

Next, we compared the pixel-level segmentation accuracy on publicly open image datasets using the wavelet-based method and Otsu’s method. Fig. 3E shows that segmentation accuracy is above 0.85 for the wavelet-based method, and smaller than 0.4 for Otsu’s method. To further test the segmentation ability in real image datasets, we chose five zebrafish embryo images and selected sub-regions that we manually curated for ground truth data and labeled the nuclei positions using our ground truth annotation tool (Fig. S8). Both the TP rate and Precision of the wavelet method had the highest scores (Fig. 3F).

### Outline of main structure and interface of the WaveletSEG matlab program

WaveletSEG is an open-source Matlab-based imaging research program we developed that runs on Windows, Mac and Linux systems with complete Graphical User Interface (GUI) and it is outlined in Fig. 4. Users can directly import either raw image files or data files with calculated results which helps them to check results or rerun processing with a different setting. 3D segmentation and quantification or other results such as embryo topology features can be easily displayed or saved in the WaveletSEG data visualization system. Users can also create scatter plots by selecting menu options to explore the relationship between them. For the examples used herein, we show the fluorescent distributions of the BMP signaling transducer phosphorylated SMAD (PSMAD) along the embryo dorsal/ventral axis, or the nuclei size spatial distribution from the outer embryo layer inwards.

**Fig. 4.**
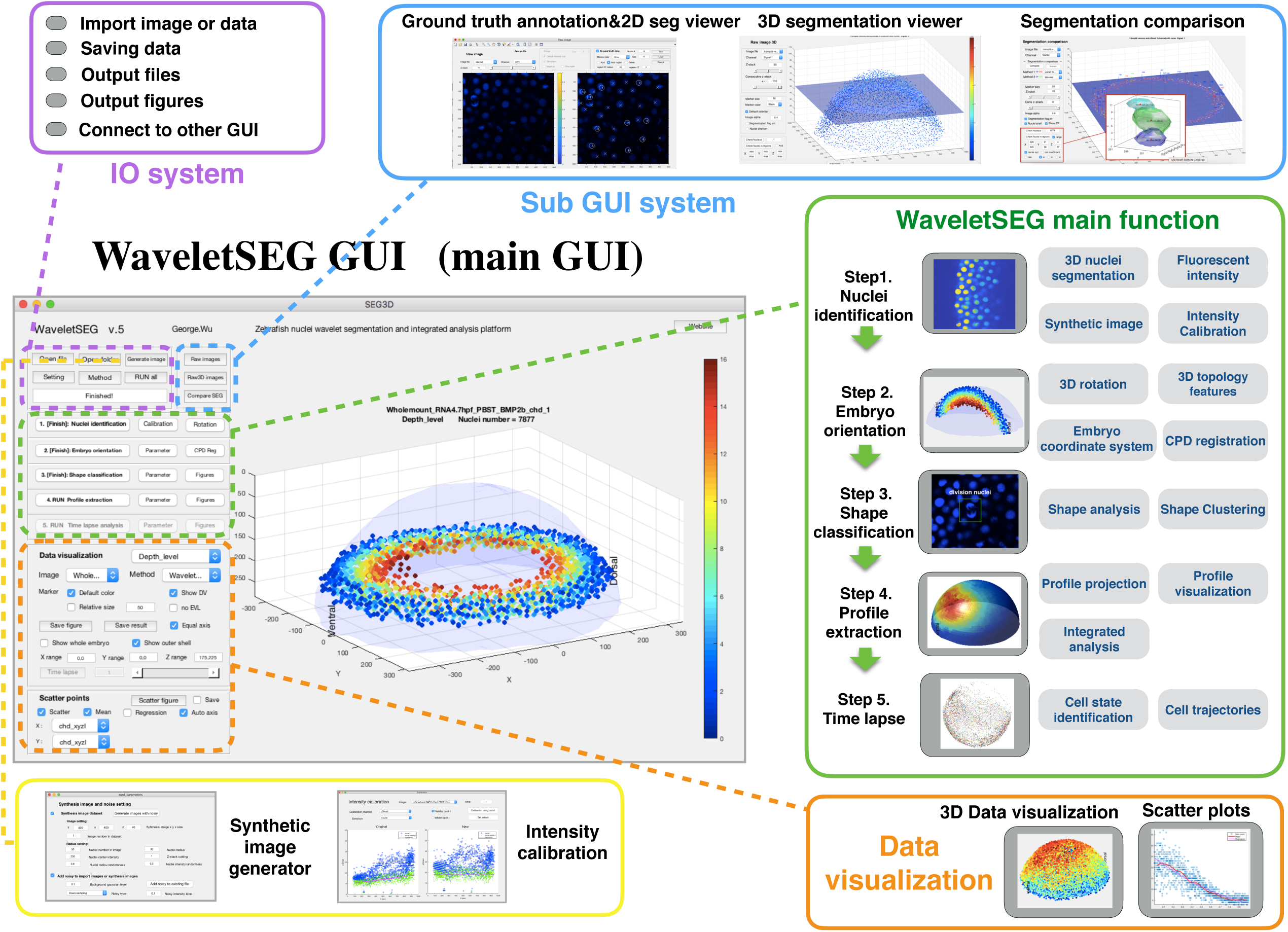
WaveletSEG software main function structure and GUI. WaveletSEG is the fully automatically user-friendly image processing analysis platform which integrated all main and extension functions including IO system, extension GUI and data visualization system in one GUI. The WaveletSEG main function block consists five main steps that can run independently including 1. Nuclei identification, 2. Embryo orientation, 3. Shape classification, 4. Profile extraction, and 5. Time lapse steps. In the nuclei identification step, users can do intensity calibration by using nuclei intensity or after wavelet-based segmentation. In IO system block, users can directly import microscope image files or intermediate data files into WaveletSEG, and save or output data results or figures directly from the GUI. We also developed a set of segmentation validation tools in WaveletSEG including synthetic data generator, 3D ground truth labeling system, 2D and 3D segmentation viewer sub-GUI, and segmentation method comparison extension GUI. In WaveletSEG data visualization block, 3D segmentation and quantification or other results such as embryo topology features can be easily displayed or saved in the WaveletSEG data visualization system. User can also create scatter plots by selecting menu options to explore the relationship between them.

WaveletSEG consists of five main steps which can run independently: 1. Nuclei identification, 2. Embryo orientation, 3. Shape classification, 4. Profile extraction, and 5. Time lapse steps (Fig. S9). In the nuclei identification step, users can do intensity calibration by using nearby nuclei intensity or do 3D rotation to adjust the embryo DV and AP axis direction after wavelet-based segmentation. After the nuclei segmentation step, the embryo coordinate system is automatically created and each nucleus is given a coordinate list value (see Methods), and the 3D topology feature for each nucleus is also calculated. In step 3, we can apply nuclei shape classifications to analyze the 3D nuclei shape type and features which resulted from step 1 and 2. In the profile extraction step, we project the nuclei quantification and features into one averaged distributed sphere plane to enhance accuracy distribution (see Online Methods). If an imported image is a 4D (3D + time) image, the user can run time-lapse analysis in step 5. In step 5, nuclei tracking is performed by segmenting all the nuclei in a time-lapse image sequence using wavelet-based segmentation. This is achieved one step at a time, and the spatiotemporal overlap between corresponding segmented regions in consecutive time steps is compared, providing information about coordinate values, topology features and shape classification.

WaveletSEG also includes a synthetic data generator, a 3D ground truth labeling system, 2D and 3D segmentation viewer sub-GUI, and segmentation method comparison sub-GUI, (see Online Methods and Fig. S7-11). Using the CompareSEG sub-GUI, users can also easily validate segmentation results by utilizing different methods and display TF, NP, or NF nuclei directly. In WaveletSEG, user can also export 2D or 3D segmentation masks as the machine learning training datasets for further object detection or semantic segmentation studies.

### Zebrafish embryo quantification

Development of an animal embryo involves molecular signaling to coordinate cellular rearrangement and proliferation. To quantify these processes in zebrafish, we quantified zebrafish BMP signaling along the AP and DV axes and calculate embryo properties including thickness, and cell division state. These properties provide a complete embryo picture (Fig. 5A) that includes the 3D embryo geometry, quantitative signaling information and other properties (Li *et al*., 2019)(Huang and Umulis, 2019). After segmentation, 3D rotation and deletion of the EVL cell later, (see Online Methods and Fig. S12-13), we generated 2D hemispherical surfaces and projected every nuclei onto the 2D surface to obtain intensity distributions of nuclear properties (Fig, 5B-C). Other outputs include whole embryo, sub region quantification results (Fig. 5D), DV axis pSMAD intensity distributions (Fig. 5G), or population average properties after the coherent point drift (CPD) registration (Fig. 5E) (see Online Methods).

**Fig. 5.**
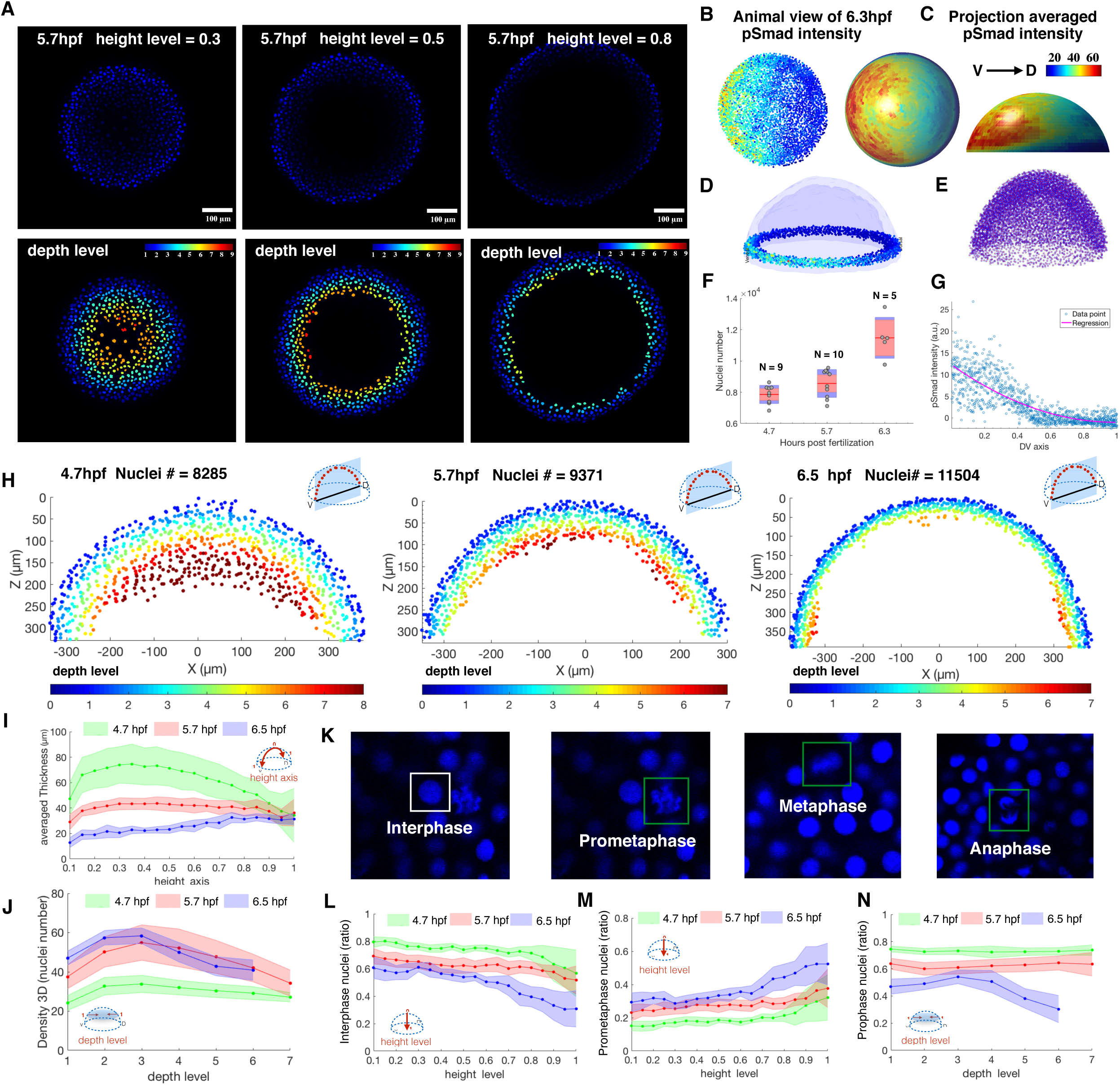
Example quantification of nuclear pSMAD gradient in Zebrafish embryo and spatial analysis of embryo coordinate system, 3D topology features and cell cycle phase patterns between 4.7, 5.7, 6.5 hpf Zebrafish embryo. (A) Z-slice DAPI nuclear stain (upper) and depth level for each nucleus on height level=0.3, 0.5, 0.8 in 5.7 hpf Zebrafish embryo (lower). (B, C) Animal view of nuclear pSMAD intensity of all nuclei from the 6.3hpf embryo (left), and animal and lateral view of averaged pSMAD intensity by projecting to uniform distributed semisphere surface using WaveletSEG (center and right). (D) Data visualization of nuclear pSMAD intensity for 6.5 hpf Zebrafish embryo in marginal region using WaveletSEG. Light blue and dark blue transparent shells indicate inner and outer boundary surfaces of all nuclei in embryo. (E) Embryo alignment using coherent point drift (CPD) method in WaveletSEG. (F) Nuclei number counts of embryo vs different developmental time (4.7hpf: N=9, 5.7hpf: N=10, 6.5hpf: N=5) which also applied in (i-j, l-n). Gray dots are individual embryo nuclei number. Red line red box region, and blue region show the mean nuclei number, 95% confidence region and region between one standard deviation region. (G) Distribution of nuclear pSMAD intensity vs DV axis in embryo margin region in (C). Blue dot and red line indicate individual nuclei intensity and averaged intensity along DV axis. (H) Depth level spatial distribution for nuclei in DV direction cross section plane ± 30 um of 4.7 hpf (left), 5.7 hpf (center) and 6.5hpf Zebrafish embryo (right). (I) Averaged nuclear layer thickness through embryo height axis in 4.7, 5.7 and 6.5hpf embryo developmental time. (J) Averaged nuclear density through embryo height axis in 4.7, 5.7 and 6.5hpf embryo developmental time. Green, red and blue dark line indicate mean value, and light green, light red and light blue regions indicate one standard deviation region. (K) Examples of cell cycle phase in mitosis based on nuclei shape classification results. (L) Prophase nuclei count ratios through embryo height level in 4.7, 5.7 and 6.5 embryo developmental time. (M) Prometaphase nuclei count ratios through embryo height level in 4.7, 5.7 and 6.5 embryo developmental time. (N) Prophase nuclei count ratios through embryo depth level in 4.7, 5.7 and 6.5 embryo developmental time. Green, red and blue dark lines indicate mean value, and light green, light red and light blue regions indicate one standard deviation region.

In early embryo development of the zebrafish, cell rearrangements guide the establishment of embryonic axes and layers(Lepage and Bruce, 2010)(Morita *et al*., 2017) and embryonic cells spread over the yolk mass while the blastoderm thins. The radial intercalation movement between deep cell layer (DCL) and enveloping layer (EVL) is regarded as the main driving force for epiboly, and the cell density increase near the EVL plays an important role in this process(Concha and Adams, 1998). Fig. 5H shows the Depth level spatial distribution for nuclei in the DV direction cross section of 4.7 hpf (N=9, left), 5.7 hpf (N=10, center) and 6.5hpf Zebrafish embryo (N=8, right) (Fig. 5F). They have significant differences not only in nuclei number, but also in the nuclei spatial distribution, density and thickness. In Fig. 5H (right), we observed the cell accumulation (Hypoblast) and involution movement near the blastoderm margin.

Emboly is another principle coordinated cell movement to form hypoblast which contains the involution and intercalation movements in the early gastrulation stage. Involuting cells move and accumulate in the blastoderm margin to narrow and elongate the embryonic axis, eventually forming endoderm and mesoderm that characterizes the gastrulation stage in Zebrafish(Bensch *et al*., 2013)(Warga and Kimmel, 1990). Convergence and extension of epiblast deep cells result from intercellular space decrease and cell density contributions to the the internalization of hypoblast cells from the margin. For further investigation of spatial and temporal analysis of embryonic structure distribution, we introduced eight 3D topology features including Size, Thickness, Density2D, Density3D, Neighbor2D, Neighbor3D, H sorting, EVL (see Online Methods and Fig. S14) which can be directly calculated in WaveletSEG. For example, Fig. 5I can be used to describe the averaged thickness in epiboly process in 4.7, 5.7 and 6.5 embryo developmental time. We also found that the thickness increases near the margin position in 6.5hpf, which is the signal for emboly process. The gradual separation of the EVL lineage correlates with the flattening of the blastoderm on the yolk cell and is accompanied by both an increase in tension and cell shape changes within the EVL(Bruce, 2016). It has been suggested that the separation of the EVL as a lineage might be the consequence of increased tension that causes EVL cell divisions to occur preferentially within the plane of the EVL. At the end of gastrula stage, the cell density is decreased at the ventral side of embryo.

Lastly, we used WaveletSEG to monitor the cell proliferation and division patterns during morphogenesis to regulate the embryo shape and growth. Nuclei segmentation and shape classification results have been used to estimate cell proliferation and cell cycle phases(Gul-Mohammed *et al*., 2014) because nuclei are spherical during early prophase, irregular shape with nuclear envelope break down, and become ellipsoidal shape in metaphase(Chan, Heisenberg and Hiiragi, 2017). Here we created the cell-cycle phases pattern in zebrafish embryo based on nuclei shape features (see Online Methods and Fig. S15).

Fig. 5M and 5N show the prophase and prometaphase nuclei count ratios at 4.7, 5.7, and 6.5hpf. We found that in all time stages, the ratio of prophase nuclei decreases from the top of the embryo to the bottom, and the prometaphase nuclei ratio increases from between 0.2 and 0.3 at 4.7 hpf to between 0.3 and 0.5 at 6.5 hpf. It showed that cell proliferation rates are low in the animal region of the embryo and higher in the zebrafish margin. These results are consistent with previous data(Mendieta-Serrano *et al*., 2013) that shows abundant mitotic cells at 50% epiboly and especially during the shield stage.

## Discussion

As illustrated above, we have developed a novel wavelet-based image segmentation algorithm for robust 3D nuclei segmentation that demonstrates good segmentation performance in many image datasets. Recent technological advances in biological imaging are focused on developing fully automated and large-scale imaging algorithms, and wavelet-based segmentation will be a good fit because of its reliability in the face of noise and intensity attenuation and the only necessary parameter is the wavelet scale factor which is chosen based on property object size and no preprocessing or fine-tuning steps are needed. We also presented a new way to divide overlapping nuclei according to the self-similarity between multiple wavelet coefficient maps.

We developed WaveletSEG to determine embryonic patterning information for molecular signaling, gene regulation, and 3D positional data for nuclei at various stages of the cell cycle. This integrated image analysis platform helps define the embryo coordinate systems to provide embryo axes for geometry features and quantification. After 3D nuclei segmentation for whole embryos, the post processing in WaveletSEG provides convenient tools to rotate the embryo, remove EVL layer nuclei and select subregions for additional quantification. Overall, this quantification and post-processing program is designed to support converting imaging data into information that can be used to infer biological mechanism, gain quantitative insight into development and develop quantitative data sets of pattern formation and signal regulation by morphogens.

## Materials and Methods

### 2D wavelet transform

1D discrete wavelet transforms (1D DWTs) and 1D continuous wavelet transforms (1D CWTs) are the most commonly applied wavelet tools in 1D signal processing and time series analysis, and are used in signal decomposition in both time domain and frequency domain(Liò, 2003)(Du, Kibbe and Lin, 2006). The 1D CWT lies in decomposition in discrete frequency domain and continuous time domain, which is different from 1D DWT. Assuming that *f*(*t*) is a continuous and integrable time-frequency signal, the 1D continuous wavelet transform (1D CWT) for *f*(*t*) can be represented by the following integral formula (equation (4),(5)):

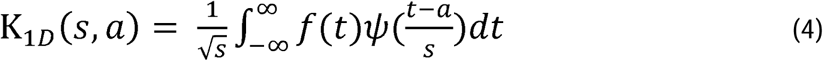

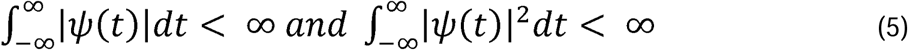

Where *ψ* /(*t*) is the wavelet mother function which must satisfy the continuous and differentiable condition. Commonly used wavelet mother functions include the Mexican hat function, Meyer function, and Morlet function. a and b are the wavelet translation factor and scale factors that represent the location shift of the mother wavelet function through time, and the dilation ratio of the mother wavelet. The output of the CWT is the wavelet coefficient matrix that is the wavelet coefficient value K_lD_ (*s, a*) in a specific translation factor a (horizontal axis) and scale factor b (vertical axis), that is also called a daughter wavelet subset. We extend the 1D CWT to the two-dimensional continuous wavelet transform (2D CWT) and apply it to the spatial domain of the image(Antoine *et al*., 1993). Supposing that *f*(*x, y*) is continuous and differentiable 2D image data, we choose the 2D Mexican hat function as the wavelet mother function and perform 2D CWT with the translation factors a and b and the wavelet scale factor s. ψ (*x, y*) is the 2D Mexican hat function and K_2D_(*s, a, b*) is the wavelet coefficient matrix (equation (6),(7)):

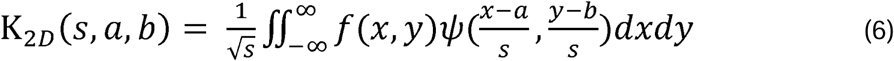

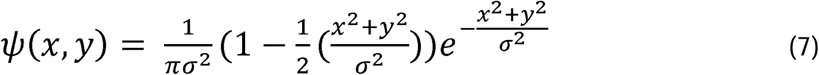

Fig. 2 shows the nuclei fluorescence microscopy raw image (Fig. 2A) and three wavelet coefficient matrices with different scale factors after the 2D continuous wavelet transform. Following application of the wavelet function (Fig. 2B), an efficient search algorithm that we developed compares different scales of CWT coefficients and marks shape-similar regions as nuclei masks in nuclei position(s) by looking for several contour lines in the CWT coefficients diagram. The similarities between maps are then calculated to determine the presence of a nuclei.

### First and second division on XY wavelet coefficient plane, XZ and YZ plane and delete small object

For this purpose, we designed division steps in two stages to divide overlapping nuclei or overlapping 2D masks using wavelet coefficient maps. For each 3D nuclei object that we obtained from the previous step, we compared Z-slice wavelet coefficient planes and found the center of each plane that is at a local minimum (Fig. 2E). If there is more than one wavelet coefficient local minimum or the displacement of center is bigger than the nuclei radius on neighboring Z slices wavelet coefficients, we assigned a new nuclei object by cutting the median line between two centers, or a new nuclei in the case of center displacement. Using the center of Z-plane wavelet coefficients, most y or z direction overlapping nuclei can be divided.

In the second-stage division step, we divided z-direction overlapping nuclei by checking both the xz plane and the yz plane wavelet coefficient (Fig. 2F). If there were more than local minimum points present in both xz and yz plane, a wavelet coefficient division plane was set in the center plane between the two centers, and a new 3D nuclei object is created. In the last step of the wavelet-based segmentation protocol, we removed the 3D nuclei voxels where the size was smaller than nuclei size × 0.2 (Fig. 2G).

### Synthetic image testing

To provide a dataset of synthetic images in validating segmentation methods, we developed a convenient interface to generate a synthetic image. Users can define synthetic image’s 3D size and the number of image and synthetic nuclei signals based on the nuclei number, nuclei radius and intensity, and the randomness of the nuclei radius and intensity between 0 to 1. If the radius randomness is 0.2, this means the generated nuclei radius = defined nuclei radius r ± uniform (0, 1) random variable × 0.2 × r. Nuclei are generated in the following order: Synthetic nuclei with random 3D positions are generated inside the image. If the distance of this nuclei relative to the previously generated nuclei, or the distance to the image boundary is smaller than nuclei r × 1.5 we regenerate the synthetic nuclei. A 3D Gaussian function with nuclei intensity and standard deviation equal to nuclei radius r × 0.4 is added to the synthetic image. We also created a synthetic overlapping nuclei dataset to evaluate the segmentation method’s ability to isolate overlapping nuclei. Subsequently, half of the nuclei were randomly created, and additional nuclei were iteratively generated and added to the dataset once its distance from the previous nuclei set was smaller than a nuclei radius r × 1.5. After initial nuclei are distributed throughout the image, we add both white noise and user-defined noise types with user-defined noise level noise density (ND) onto the synthetic image that we generated in the previous step. White noise is white Gaussian noise with user-defined variance. User-defined noise types include Gaussian white noise and salt-and-pepper noise with ND and intensity attenuations with image gradient ND, in which pixel intensity will increase continuously from right to left with intensity ND times on the left side of the image.

### Point-wise method, Otsu method, DS method and parameter screening using GA

To evaluate the segmentation performance of the wavelet-based segmentation method, we compared it with three commonly used segmentation methods, including the point-wise method, the Otsu method and the DS algorithm. The point-wise method is the simplest method to estimate the intensity of nuclei center points i. Raw images were firstly smoothed using 9 × 9 × 3 kernel, followed by h-maxima and h-minima transform to suppress all background signals.

After combining nearby local maxima closer than 6 pixels, the remaining local maxima were assumed to be nuclei center points and intensity was calculated after applying 6×6×3 spherical kernel. The Otsu method is the most popular threshold-based segmentation method to separate foreground and background pixels by finding the optimum threshold to satisfy minimum intra-class variance. We applied a symmetric Gaussian low-pass filter and sharpened the filter with specific radius and amount on the raw image. We then used Wiener filter or pixel-wise adaptive low-pass Wiener filter to deblur the image and remove noise. To this end, we used the multilevel thresholding Otsu method to decide threshold values.

Derivatives Sum algorithm (DS algorithm) started with applying denoising filters such as a Gaussian filter or a non-linear isotropic diffusion filter on the image, and the 2D spatial derivatives were computed to get the image Gauss gradient, Laplacian determinant, and Hessian determinant. A mask function F was calculated by combining the first and second spatial derivatives with weight parameters, and the Otsu method was applied here to obtain binary image slices. 3D nuclei segmentation was obtained by connecting the neighboring 2D masks, and we used a genetic algorithm for the parameter screening in this method(Bäck and Schwefel, 1993).

### Ground truth labeling GUI

Creating embryonic 3D nuclei segmentation ground truth data is challenging and time-consuming. Here we introduced Raw_image sub-GUI, a 3D nuclei annotation tool that provides a convenient and efficient way to label 3D nuclei, which can also be used to evaluate nuclei segmentation results.

In Raw_image the user can directly label embryo nuclei by simple mouse clicking. There are two main types of labeling: nuclei center labeling that shows the nuclei index, and non-center nuclei labeling that is marked as a cross symbol. Non-center nuclei labeling can help to prevent double counting. After clicking on the screen, one sub-GUI will jump up with the specific region and a range of Z-slack planes (Fig. S8B). The user can directly assign the labeling by clicking. Raw_image can also be used to check segmentation results by either comparing segmentation masks with raw images or checking single nuclei masks.

### CPD registration method

The coherent point drift (CPD) algorithm is a probabilistic-based point set registration method widely used in the field of pattern recognition(Myronenko and Song, 2010). The main goal of point registration is to merge two or more points sets into one representative set. CPD treats the registration process as a probability density estimation problem and fits the Gaussian mixture model (GMM) of one-point set to a reference point set using maximum likelihood method, and the expectation maximization algorithm is used to make the most effective use of the optimization function. We use CPD to merge embryos in the same stage into a representative embryo one by one. The final GMM centroids are the representative embryo nuclei sets.

### 3D topology features

We designed eight 3D topological features to describe the topology and geometric structural changes for embryological development studies in Zebrafish. Nuclei size is the total pixel numbers for every 3D segmented nuclei. We used four indexes to describe the spatial nuclei density distribution for the embryo, including density 3D, density 2D, neighbor 3D and neighbor 2D distance. Density 3D is the total nuclei number with distance smaller than 30 pixels around one nucleus. The problem of using density 3D is the underestimation near the outermost or innermost layer. In this instance, we define density 2D as the total nuclei number closer within 30 pixels distance and also in the same layer (depth level). Neighbor distance is the average distance of the nuclei to the closest two nuclei which is suitable to find the outlier point in point set. The difference between neighbor 2D and neighbor only considers nuclei in the same layer.

Another important topological feature of the embryo is the thickness. We define thickness here as the vertical distance from the innermost layer of embryo. The innermost shell was divided into 360 sub-regions, and the vertical distance was calculated from nucleus to its corresponding plane. The outer epithelial monolayer of a zebrafish embryo is called the enveloping layer (EVL). Here we defined the inner EVL and outlier EVL as the layer with depth level equal to one or the maximum values. H_sorting is the index to describe the dynamic process of nuclei movement by dividing every 500 nuclei from the top of the embryo to the embryo button.

### Cell cycle phase pattern based on nuclei shape classification

To study the spatial and temporal shape distribution of embryo nuclei, we defined some rules relative to dividing them into spherical nuclei, irregular shape nuclei, elliptical nuclei and dividing nuclei. We calculated the sphericity value using 4 × pi × area divided by squared perimeter in three planes (xy, yz and xz planes), and averaged them as the averaged sphericity value. The average aspect ratio is the mean of aspect ratio xy, yz, and xz. Aspect ratio xy is the proportional relationship between x axis range divided by y axis range.

Here we defined spherical nuclei if the mean sphericity value is larger than 0.9 which means the shape of nuclei is similar to a circle with the same area. If the mean sphericity value is smaller than 0.9 and the mean aspect ratio is larger than 0.7, we identified it as an irregular shape of nuclei. A nucleus is determined as either elliptical or dividing if the mean sphericity value is smaller than 0.9 and the mean aspect ratio is also smaller than 0.7. In addition, if the radius of this nuclei is smaller than the embryo mean radius and they are paired, they are considered dividing nuclei.

### Embryo profile projection and data visualization

To indicate a better concentration or feature distribution on a three-dimensional multi-layer embryo, and also allow for the integration of similar stage embryos, we projected features of each nucleus into one reference two-dimensional surface. We generated a hemispherical surface with averaged distributed reference point which we created in using regular placement method. We projected the nuclei into reference points with the closest spherical coordination system angle (θ, φ). Thereafter, we summarized the concentration of nuclei in this reference point, as well as average topological features such as nuclei size, nuclei density or thickness.

We designed the interactive graphical user interface to visualize segmentation and analyze the results of the embryo directly. Users can also choose a particular region such as the embryo margin region by choosing z region range, or DV axis concentration distribution. In addition, users can remove EVL nuclei directly by clicking the no EVL icon on the data visualization board. In a scatter plot board, users can choose one coordinate system index in the first pop-up menu and one feature in second pop-up menu to analyze the spatial distribution on the embryo. Alternatively, users can also plot the relationship between two features using the scatterplot board.

### Zebrafish lines

All the procedures done on zebrafish adults and embryos were approved by the Purdue University Institutional Animal Care and Use Committee (IACUC). Zebrafish were raised and maintained by the Pentair Aquatic Eco-Systems. Wide-type (TL) embryos were collected in E3 water when a one male-and-female pair was crossed for 30 min. Subsequently, embryos were left to develop to the desired stage at 28 °C.

### Immunostaining

Embryos were penetrated in 0.1% Triton X-100 in PBS for 1 hour at RT. Embryos were blocked in blocking buffer (4% BSA(Millipore Sigma, #126615), 1% DMSO, 0.1% Triton X-100 in PBS) overnight at 4°C, and then stained with anti-phosphoSmad1/5/8 antibody (Cell Signaling Technology, #9511) at 1:100 diluted in blocking buffer overnight at 4°C. Then embryos were detected by goat anti-rabbit cross-adsorbed Alexa Fluor 647-conjugated antibody at 1:500 dilution with DAPI overnight at 4°C (Thermo Fisher Scientific, #A21244).

## Code availability

WaveletSEG is freely available for academic use as Supplementary Software or (including updated versions) at https://waveletseg.weebly.com/. Source code is available upon signing of a material transfer agreement.

## ACKNOWLEDGEMENTS

We thank the help of Matt Thompson and Michelle Ingle for generating the nucleus ground truth data; Madeline Ku for codes testing.

## Data availability

Raw data and segmentation result files that support the findings of the study are available at https://waveletseg.weebly.com/.

---

**Fig. S1. Illustration of wavelet-based segmentation method in an optical section of a whole-mount zebrafish embryo**. (A, A1-3) Whole-mount embryo z-slice nuclear DAPI-staining raw images of germ ring-stage (5.7 hpf) zebrafish and three red-boxed regions. Image data were acquired with a Zeiss LSM 800 confocal microscope with 20X water lens. (B, B1-3) Wavelet coefficient matrixes after applying 2D CWT on raw images in (A). The blue color corresponds to the negative coefficient value region, and the yellow color corresponds to the positive coefficient value region. 2D Mexican hat function was selected as the mother wavelet function. (C, C1-3) 2D segmentation masks (yellow regions) were obtained after applying multi-scale object identification on wavelet coefficient matrixes in (B).

**Fig. S2. Limitations for nuclei segmentation on whole embryo confocal imaging**. (A, B) Nuclei with clear boundaries high contrast and low contrast and non-uniform background intensity nuclei. (C) Whole zebrafish embryo z-slice nuclear DAPI staining with confocal Intensity attenuation effect, and the nuclei segmentation results using wavelet-based segmentation method. Green boundaries show nuclei segmentation masks and nuclei with inhomogeneous intensity can be detected.

**Fig. S3. Nuclei segmentation and RNA segmentation results using wavelet-based segmentation method**. (A) C.elegans developing embryo nuclear DAPI staining by light-sheet microscope with segmentation masks marked by green boundaries (B) Pancreatic Stem Cells on a Polystyrene substrate, (C) GFP-GOWT1 mouse stem cells, and (D)(E) Whole mount zebrafish embryo z-slice bmp2b mRNA expression at 5.7hpf, and bmp RNA spot segmentation results using wavelet-based segmentation method in different z planes(left and right). Insets: enlarged views of two while box regions.

**Fig. S4. Workflow of multi-scale object identification step in wavelet-based segmentation method**. (A) Zebrafish embryo nuclear DAPI staining raw images (5.7hpf). We apply 2D continuous wavelet transform (2D CWT) on this raw image with three wavelet scale factors s1, s2, s3. (B) Three wavelet coefficient maps with wavelet scale factors s1, s2, s3. Contour color represents intensity of the corresponding wavelet coefficient. The green intensive contoured ridges region in three wavelet coefficient maps are almost identified and can be correlated with nuclei region in raw image. (C) Local minimums were determined on three wavelet coefficients map after the H-minima transform. If local minimums appeared in the same positions on all wavelet coefficient maps, they were identified as potential nuclei peak locations and the corresponding zero-value cross sections are calculated. (D) In this step we calculated the 2D masks intersection over Union (IOU) of all zero-value cross sections from all wavelet coefficient maps. If the difference in cross section areas is less than 10%, they averaged and were identified as nuclei 2D masks. (E) The final segmentation mask after multi-scale object identification step.

**Fig. S5. First division step based on wavelet coefficient center positions on z-slice of nuclei 3D blob**. (A) Five z-slice wavelet coefficient maps of segmented nuclei 3D blob from top (z = 7) to bottom (z = -1). Color represents intensity of the corresponding wavelet coefficient from positive value (yellow) to negative value (deep blue). Red cross marks the center position (local minimum position) on each wavelet coefficient map z-slice in 3D blob. Here the z value came from z axis of whole embryo before 3D rotation. (B) If there are more than one center positions found or the shift of the center positions between neighboring Z slices is bigger than nuclei radius, we assigned a new nuclei object by cutting the median line between two centers, or assigned new nuclei in the case of center displacement. In (A) we found two center position shift between z=5 to z=3 and z=1 to z=-1, and marked using red arrow. So we divided the original 3D nuclei blob (left) into three nuclei 3D blobs (right) with three new nuclei centers (red dot) and displayed using different color. (C) Three examples show the 3D nuclei blobs after the first division step. Red box on the third 3D nuclei blob (right) shows the nuclei overlapping in z direction cannot be divided in the first division step.

**Fig. S6. Second division step in z direction and the summary of blob size distribution before and after division steps**. (A) The 3D nuclei blob (left) shows the nuclei overlapping in z direction which didn’t divide appropriately in the first division step because there is no center position shift between neighboring Z slices. In the second division step, wavelet coefficient yz and xz plane are exanimated and search for center positions for every 3D nuclei. If two center positions are found on both yz and xz planes (center left and center right), original 3D nuclei blob will be divided into two new blobs from the middle plane of two center positions (right). (B) Remove the 3D nuclei blobs smaller then nuclei size × 0.2. (C) Blob size distributions in three axis directions (upper: x-direction, center: y-direction, lower: z-direction) before the first division step (left column), and after the first division step (center column), and after the second division step (right column) for all nuclei segmented using wavelet-based segmentation method in whole embryo. The final size distributions in three directions are all similar to normal distributions (right column).

**Fig. S7. The workflow of synthetic data generator and process to add noise in WaveletSEG**. The synthetic image generator in WaveletSEG can generate synthetic images or synthesis overlapping images by providing image number, image size, synthetic nuclei number, nuclei radius and intensity and their randomness. For overlapping images, we iteratively create pairing nuclei with distances between two nuclei are smaller than nuclei radius × 1.5. After initial synthesis image dataset are created, Gaussian noise, salt and pepper noise with different noise level can be added into images, or apply down-sampling or up-sampling to those images.

**Fig. S8. Ground truth image dataset we used and the ground truth labeling GUI and 2D segmentation viewer**. (A) Five 3D ground truth of 400×400×24 voxel region nuclear DAPI staining raw images from 4.7hpf Zebrafish embryo (first three images), and raw images 400×400×24 voxel region of 4.7hpf Zebrafish embryo in pSMAD datasets. (B) To build the ground truth dataset using our own images to validate the segmentation method, we introduced Raw_image sub-GUI, a 3D nuclei annotation tool which provides a convenient and efficient way to label 3D nuclei, and can also be used to evaluate nuclei segmentation results. The left window of the Raw_image shows the raw image with side bar to view different z-slice. (C) The right window shows both raw image and segmentation labels, and provides a very convenient way to label 3D nuclei. When user clicking inside the right side window, pop-up GUI will show up and display every z-slice in this position. User can very easy to select the nuclei center and do labeling.

**Fig. S9. WaveletSEG main functions**. WaveletSEG main function block consists of five main steps which can run independently including 1. Nuclei identification, 2. Embryo orientation, 3. Shape classification, 4. Profile extraction, and 5. Time lapse steps. In the nuclei identification step, whole embryo nuclei are segmented using wavelet-based segmentation method, and averaged nuclear fluorescent intensity is also calculated. User can do intensity calibration by using nuclei nearby intensity or do 3D after wavelet-based segmentation. In IO system block, User can directly import microscope image files or intermediate data files into WaveletSEG, and save or output data results or figures directly from the GUI. In the embryo orientation step, individual embryo is rotated in 3D automatically according to some specific chemical such as chd mRNA. Embryo coordinate value and 3D topology features for each nucleus is also calculated after rotation to provide positional information. To integrate multiple embryos, CPD registration is applied here and created one representative embryo. In shape classification step, we defined shape-based rules to classify nuclei into spherical nuclei, irregular shape nuclei, elliptical nuclei, and division nuclei to estimate cell proliferation and cell cycle phases. Unsupervised learning clustering such as SOM network is also included to cluster nuclei based on nuclei size and shape. In profile extraction step, we project the nuclei quantification and features into one averaged distributed sphere plane to enhance accuracy distribution. We also developed a set of segmentation validation tools in WaveletSEG including synthetic data generator, 3D ground truth labeling system, 2D and 3D segmentation viewer sub-GUI, and segmentation method comparison extension GUI. In WaveletSEG data visualization block, 3D segmentation and quantification or other results such as embryo topology features can be easily displayed or saved in WaveletSEG data visualization system. User can also create scatter plots by selecting menu options to explore the relationship between them.

**Fig. S10. Raw image3D sub-GUI**. Raw image3D is the 3D segmentation data visualization GUI to check and validate the segmentation results in WaveletSEG, and user can execute this sub-GUI by clicking the button “Raw3D images”. Blue points inside the main data visualization window (Right) are segmented nuclei positions are displayed as 3D point cloud, and horizontal plane is the raw image z-slice enable user to validate and compare 3D segmented nuclei with original raw image z-slice. Left side of this sub-GUI is the control panel to display regional nuclei or change into different images or channels.

**Fig. S11. Compare SEG sub-GUI**. Compare SEG is designed to validate the segmentation results or compare segmentation results using different segmentation methods in WaveletSEG. User can execute this sub-GUI by clicking the button “Compare SEG” in WaveletSEG. Right side of this sub-GUI is the main data visualization window and segmented nuclei positions are displayed as 3D point cloud.

**Fig. S12. Zebrafish embryo coordinate system in WaveletSEG**. we developed the zebrafish embryo coordinate system including embryonic axes such as DV axis, AP axis, and DV center axis, Height axis, Height level axis, Depth level axis. If time-lapse images are available, there is an additional axis: time stage. The first five coordinates are values between zero to one, Depth level is the layer number counting from the outer shell, and the Time stage started from 1. DV center axis is defined by connecting ventral point to dorsal point through the top of embryo, angle is calculatedF and DV center value is from 0 to 1. Unlike the Height level axis which described the vertical distance from top to button of Embryo, we defined Height axis through the line connecting ventral point to dorsal point through the top of embryo to describe the embryo developmental process relative to which nuclei will grow from the top of the embryo to the embryo bottom.

**Fig. S13. Data workflow from 3D whole embryo raw image to projected average intensity distribution in WaveletSEG**. Step1: Import 3D raw image with both DAPI stained channel and pSMAD channel. Step2: Segmented nuclei point cloud by using wavelet-based segmentation method. Step3: Automatically rotation of embryo by using chd or pSMAD channel embryonic spatial distribution. Step3a (optional): Apply intensity calibration by using nearby nuclei intensity or do 3D rotation to adjust the embryo DV and AP axis in the right direction after wavelet-based segmentation. Step3b (optional): To merge embryos in the same stage into a representative embryo one by one using CPD. Step4: After 3D rotation and embryo center point is moved to position zero. Step5: Automatically remove the outermost layer and the innermost layer from embryo using depth level information. Step5a (optional): data analysis and data visualization in specific sub-region of whole embryo. Step6: We generated a hemispherical surface with averaged distributed reference points using regular placement method, and projected each nucleus into this 2D hemispherical surface (projection surface). Step7: We projected the nuclei into reference points with the closest spherical coordination system angle (θ, φ) and created a projected average intensity and 3D topology.

**Fig. S14. 3D topology features used in WaveletSEG**. Eight 3D topology features are defined in WaveletSEG software to quantitatively describe topology changing through embryo developing process. Nuclei size is the total pixel number for every 3D segmented nuclei. We used four indexes to describe the spatial nuclei density distribution for the embryo, including density 3D, density 2D, neighbor 3D and neighbor 2D distance. Density is the total nuclei number with distance smaller than 30 pixels around one nucleus, and Neighbor distance is the average distance of the nuclei to the closest two nuclei. The difference between 3D and 2D is that we only calculate density in the same layer (depth level) for density 2D and neighbor 2D distance. Thickness is defined as the vertical distance from the innermost layer of embryo. The outer epithelial monolayer of a zebrafish embryo is called the enveloping layer (EVL). Here we defined the inner EVL and outlier EVL as the layer with depth level equal to one or the maximum values. H_sorting is the index to describe the dynamic process of nuclei movement by dividing every 500 nuclei from the top of the embryo to the embryo bottom.

**Fig. S15. Cell cycle phase patterns based on nuclei shape classification results**. (A) Examples of cell cycle phase in mitosis based on nuclei shape classification results, and the defined shape-based rules for cell cycle phase. (B) Nuclei count ratios for prophase, prometaphase, metaphase, and anaphase nuclei along the embryo height level in 4.7, 5.7, and 6.5 embryo developmental time. (C) Nuclei count ratios for prophase, prometaphase, metaphase, and anaphase nuclei along the embryo depth level in 4.7, 5.7, and 6.5 embryo developmental time. Green, red, and dark blue lines indicate mean value, and light green, light red, and light blue regions indicate one standard deviation region.

